# The role of auxin-mediated gene activation in the bryophyte, *Physcomitrium patens*

**DOI:** 10.1101/2024.10.15.618431

**Authors:** Carlisle Bascom, Danielle Tu, Mark Estelle

## Abstract

Perception and response to the hormone auxin is critical to plant growth and development. Expression of auxin-response genes is tightly regulated via known mechanisms of both activation and repression. Across the plant lineage, auxin-response gene induction is performed by AUXIN-REPSONSE FACTOR (ARF) activating transcription factors. Conversely, AUXIN/INDOLE ACETIC ACID proteins repress expression. Studies of gain-of-function constitutive-repression lines and *ARF* loss-of-function mutants have advanced the field. Yet, there is a need for a comparative study of aberrant auxin-signaling mutants to understand the developmental consequences of constitutive repression versus the absence of auxin-mediated gene induction. Using CRISPR/Cas9 gene-editing tools, we mutated each activating *ARF* gene in the model bryophyte, *Physcomitrium patens.* The resulting septuple loss-of-function mutant line (*arfa^sept^*) has severe developmental phenotypes and a diminished ability to respond to exogenous auxin. However, phenotypic analysis revealed that the *arfa^sept^* line is not as severely affected as the constitutive-repression lines. Expression analysis of several auxin-response genes demonstrate that auxin-mediated gene induction is abolished in both *arfa^sept^* and constitutive-repression lines but that basal expression levels are higher in the *arfa^sept^*lines. Our results suggest that the expression of auxin-regulated genes important for developmental progression is maintained, albeit at reduced levels, in the absence of ARFs.

**Highlight:** Researchers used CRIPSR/Cas9 to produce a septuple Auxin Response Factor mutant in the model bryophyte *Physcomitrium patens* revealing the consequences of the complete loss of auxin-mediated gene activation on development.

## Introduction

Decades of research have underscored how pivotal the phytohormone auxin is to plant growth and development (Gomes and Scortecci 2021). Despite immense genome diversity across the plant lineage (Harris et al. 2022), the molecular mechanisms for nuclear auxin signaling remain conserved from bryophytes to angiosperms (Bowman et al. 2017; Flores-Sandoval, Eklund and Bowman 2015; Matthes et al. 2019; Rensing et al. 2008). Auxin-induced gene expression occurs as the result of de-repression of transcription (Salehin, Bagchi and Estelle 2015). Auxin Response Factors (ARFs) bind to the promoter region of auxin-response genes via the B3 DNA Binding Domain. However, the effect of ARFs on gene expression is limited due to their interaction with AUXIN/INDOLE ACETIC ACID (Aux/IAA) transcriptional repressors. ARF-Aux/IAA interactions occur through their mutual Phem/Box1 (PB1) domains (Guilfoyle 2015; Sumimoto, Kamakura and Ito 2007). Aux/IAA proteins recruit the transcriptional co-repressor TOPLESS (TPL) (Szemenyei, Hannon and Long 2008) which represses gene expression. The phytohormone auxin facilitates the interaction of Aux/IAA proteins and the TRANSPORT INHIBITOR RESISTANT 1/AUXIN F-BOX (TIR1/AFB) subunit of SCF^TIR1/AFB^ E3 ubiquitin ligases. This interaction results in the ubiquitination and degradation of Aux/IAA proteins, thereby alleviating transcriptional repression. In the absence of Aux/IAA proteins, the ARFs can promote transcription of auxin regulated genes.

Many studies have defined three major classes, or clades, of ARFs; activating (A), repressing (B), and the more enigmatic C ARFs (Finet et al. 2013; Li et al. 2016; Mutte et al. 2018; Tiwari, Hagen and Guilfoyle 2003). Further, recent work has highlighted the fact that bryophytes possess clade D ARFs, also called non-canonical ARFs, that function as activators despite not having a DNA Binding Domain (Bascom et al. 2023; Mutte et al. 2018). Clade D ARFs are also present in lycophytes, but to date there are no published studies on lycophyte ARFs. Among the activating ARFs, the first and best studied is *MONOTOPOROUS/AUXIN RESPONSE FACTOR 5 (MP/ARF5)* in *Arabidopsis* (Hardtke and Berleth 1998; Ulmasov et al. 1997; Wójcikowska, Belaidi and Robert 2023). Null mutants of *AtARF5* fail to produce hypocotyls, but can be grown to flowering age by inducing adventitious roots (Przemeck et al. 1996). Further, *mp/arf5* plants produce far fewer flowers than wild type as MP/ARF5 protein directly activates gene expression of the floral-fate determining gene *LEAFY* (*LFY)* (Yamaguchi et al. 2013). Later work demonstrated that MP/ARF5 protein interacts with the chromatin remodelers BRAMA (BRM) and SPLAYED (SYD) to activate gene expression (Wu et al. 2015). BRM and SYD are members of the SWItch/Sucrose Non-Fermentable (SWI/SNF) family of ATPases with profound effects on gene expression across eukaryotes (Bieluszewski et al. 2023). Importantly, BRM and SYD proteins compete with Aux/IAA proteins for binding to MP/ARF5, such that BRM/SYD-MP/ARF5 interaction is facilitated by the auxin-mediated degradation of Aux/IAA (Wu et al. 2015).

Researchers have used the bryophytes *Physcomitrium patens* (formerly *Physcomitrella,* a moss) and *Marchantia polymorpha* (a liverwort) to understand the derivation and evolution of auxin signaling (Flores-Sandoval, Eklund and Bowman 2015; Thelander, Landberg and Sundberg 2018). Recent work demonstrated that mutating all *TIR1/AFB* genes in *Arabidopsis,* thereby eliminating the canonical auxin signaling pathway, is embryo lethal (Prigge et al. 2020). In contrast both model bryophytes are able to survive without their AFBs, albeit in a severely stunted state (Bascom et al. 2023; Suzuki et al. 2022). Mutations in the degron motif, “Domain II”, of Aux/IAAs result in the repression of auxin signaling, as the IAA proteins are not degraded in response to auxin (Zenser et al. 2001). Phenotypes attributed to gain-of-function degron mutations in *Arabidopsis* are generally more severe than null alleles (Tian, Uhlir and Reed 2002). The expanded *Aux/IAA* gene family and genetic redundancy in *Arabidopsis* has made loss-of-function studies difficult. Much of what we know is based on degron mutants (Luo, Zhou and Zhang 2018). Likewise, experiments with gain-of-function *Aux/IAA* mutants in *M. polymorpha* (Kato et al. 2015) and *P. patens* (Prigge et al. 2010) reinforced the developmental consequences of constitutive repression of auxin-regulated genes. Only in *P. patens* has Aux/IAA function been completely deleted (Lavy et al. 2016). The loss of all three moss *Aux/IAA* genes results in a constitutive auxin response and severely stunted growth demonstrating that Aux/IAA-mediated gene repression is an essential part of development.

The *M. polymorpha* genome contains a single AFB and Aux/IAA gene as well as single gene from each clade of (Bowman et al. 2017). *M. polymorpha* lines lacking the single activating ARF, *MpARF1,* are unresponsive to exogenous synthetic auxin 1-naphthaleneacetic acid (NAA), and have reduced endogenous auxin signaling (Kato et al. 2017). Predictably, *mparf1* lines have severe developmental phenotypes and are stunted compared to wild type (Kato et al. 2017). In contrast to *M. polymorpha*, the *P. patens* genome contains eight activating ARFs (ARFas), four AFBs (Bascom et al. 2023; Prigge et al. 2010), and three Aux/IAAs (Lavy et al. 2016; Prigge et al. 2010). Previous work demonstrated that inducible PpARFa8 expression can restore auxin-mediated gene activation in a line where auxin response genes are repressed solely by an ectopically expressed repressing ARF (Lavy et al. 2016). Intriguingly, constitutive expression of PpARFa8 in a wild type background results in plants that superficially resemble plants expressing stabilized Aux/IAA proteins (Lavy et al. 2016). These results confirm models whereby PpARFas recruit Aux/IAAs to the promoter region of auxin-sensitive genes. To better understand the role and function of activating ARFs in plant development, there is a need to directly compare lines in which each component of the nuclear auxin signaling pathway has been disrupted separately.

Given the ease of genetic manipulation and position at the phylogenetic base of land plant evolution, *P. patens* is a well-suited to a comparative analysis of auxin signaling mutants. Like other land plants, several important developmental steps are auxin mediated (Landberg et al. 2020; Thelander, Landberg and Sundberg 2018, 2019). The *P. patens* life cycle starts as haploid spores germinate to form a filamentous tissue known as protonema. Protonemal cells consist of two morphologically and functionally distinct cell types– chloronemata and caulonemata. The apical chloronemal cell in a growing filament differentiates into a caulonemal cell in an auxin-dependent manner (Plavskin et al. 2016). Additionally, the switch to three-dimensional growth is also an auxin-mediated process. Gametophores, the “leafy” moss structures that grow against the gravity vector, initiate as buds along caulonemal cells. The initiation and development of gametophores has been identified as an auxin-mediated process that is independent from that of the chloronemal-caulonemal transition (Landberg et al. 2013). Both the formation of caulonemal cells and normal development of gametophores have been used as indicators of a fully functioning nuclear auxin signaling pathway (Eklund et al. 2010; Thelander, Landberg and Sundberg 2018; Viaene et al. 2014).

In this study we explore the function of the ARFa proteins in *P. patens* development. We mutated all seven *ARFa* genes, thus generating a line that completely lacks auxin-mediated gene activation. This is in contrast to *afb-*null and Aux/IAA*-*stabilized lines, in which the phenotypes are the result of increased or constitutive repression of auxin-regulated genes. Phenotypes of an *ARFa-null* plant reveal the developmental body plan in the absence of auxin response gene activation. Further, mutant lines devoid of any functional activating ARF, when compared to lines in which auxin signaling is repressed at other stages of the pathway, would reveal the relative contribution of PpAFBs, PpAux/IAAs, and PpARFas to auxin signaling specifically and plant development broadly. Our conclusions largely support the de-repression model of nuclear auxin signaling, but highlight the likely importance of non-ARF transcription factors on the expression of auxin-responsive genes.

## Results

### Mutation of the ARFa genes has severe consequences on gametophore development and auxin response

To understand the developmental role of PpARFas, we generated higher order *arfa* mutant lines (Figure 1A). The *P. patens* genome encodes eight *ARFa* genes (Rensing et al. 2008). The *ARFa1* (Pp3c1_14480v3) and *ARFa2* (Pp3c1_14440v3) loci are identical except for a C to T mutation in *ARFa2.* If transcribed, the C to T mutation in *ARFa2* generates a premature stop codon within the DBD. However, publicly available annotations of the *ARFa2* splice around the nonsense mutation, which has the consequence of omitting conserved regions of the DBD (Goodstein et al. 2012). As such, the 3’ portion of the fourth intron which contains the C to T mutation and entire fifth exon of *ARFa2* are identical to the sixth exon of *ARFa1.* Transcriptomic data does not support the expression of *ARFa2’s* truncated fifth exon, nor the C-terminal region where ARFa1 and ARFa2 protein sequences diverge (Goodstein et al. 2012; Lavy et al. 2016). Based on these results, we conclude that *ARFa2* is a pseudogene that is presumably the product of the duplication of *ARFa1* that subsequently developed a nonsense mutation in the DBD encoding region of the gene. As such *ARFa2* is a pseudogene and thus was not part of our study. Finally, it is likely that RNA-sequencing reads mapped to *ARFa2* are actually from *ARFa1*. For the remaining seven *ARFa* genes, we generated null mutations by successive transformations with a transient CRISPR/Cas9 system targeting the ARFas alone or in pairs (Mallett et al. 2019). After each transformation, the target loci in recovered plants were screened via PCR for insertions and/or deletions (Supplemental Figure 1). The last two ARFa genes targeted were *PpARFa1* and *PpARFa3*, and we recovered two independent lines from that transformation event (see Materials and Methods). With the exception of *arfa7-6,* each mutant allele contained clear deletions or insertions (Supplemental Figure 1) that resulted in a nonfunctional protein (Supplemental Figure 2). For the *arfa7-6* allele, sequence analysis revealed the presence of an in-frame deletion (Supplemental Figure 2). While the deletion eliminated conserved regions upstream of the DBD, the DBD itself was left intact. Therefore, we designed a new CRISPR/Cas9 construct to target the *arfa7-6* locus. The resulting *arfa7-78* allele has a portion of the 3′ Untranslated Region (UTR) inverted and inserted into the 5′ end of the coding sequence (Supplemental Figure 1). Subsequent phenotyping indicated that *arfa* septuple mutants carrying either *arfa7-6* or *arf7-78* were identical (Figure 1), and therefore *arfa7-6* is likely null. Further, a comparative meta-analysis of *ARFa* gene expression consistently showed *PpARFa7* is the least expressed of *ARFas* in protonemata (Supplemental Figure 3), and ARFa7 protein likely contributes comparatively little to the over-all nuclear auxin response in this tissue type. In conclusion, we were able to generate two *arfa^sept^* mutant lines with independent alleles of *arfa1, arfa3* and *arfa7*.

**Figure 1.**
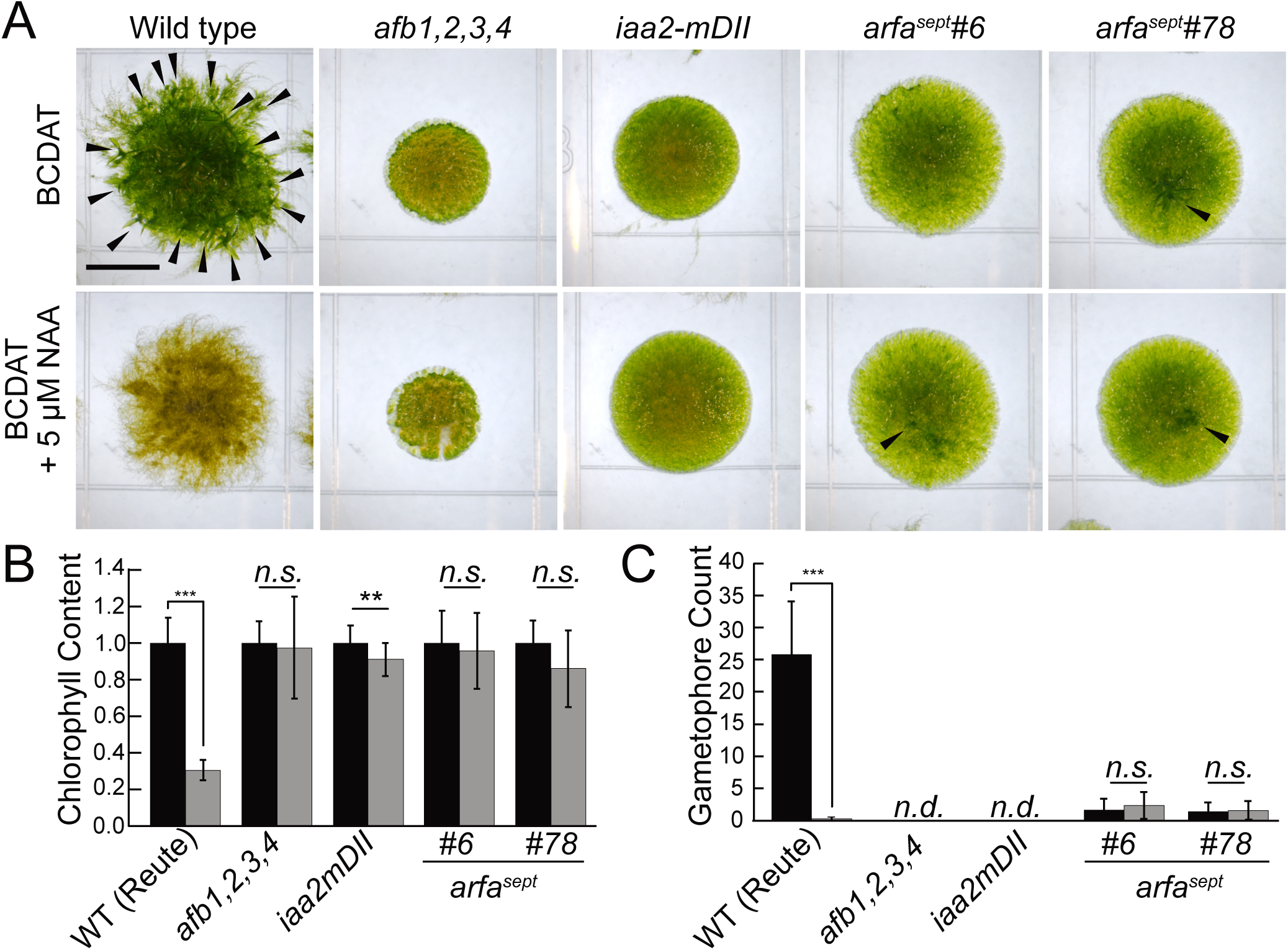
*arfa^sept^* mutants have severe developmental phenotypes. A) 21-day-old moss colonies grown on BCDAT. Each colony developed from a spot inoculum of protonemal tissue. Black arrowheads indicate select gametophores on WT and *arfa^sept^* colonies. Scale bar 5.0 mm. B) Average chlorophyll content of 21-day-old moss colonies grown on BCDAT ± 5.0 µM NAA normalized to mock-treatment. Error bars are standard deviation. *n* ≥ 11 colonies across two replicates. C) Average number of leafy gametophores that colonies developed after 21-days on BCDAT ± 5.0 µM NAA. Error bars are standard deviation. *n* ≥ 18 colonies across three replicates.

To observe developmental phenotypes in the *arfa^sept^* lines, we compared 21-day-old colonies grown from the spot inoculations of equal amounts of protonemal tissue of wild type and mutant lines (Figure 1). After three weeks, wild-type colonies are richly green, the peripheries are uneven due to the development of caulonemal filaments, and dozens of gametophores are clearly visible (Figure 1A). While both *arfa^sept^* colonies are also green, the colonies have a smooth periphery and few gametophores develop. Given that the development of caulonemata and gametophores is auxin-mediated, it is clear that the auxin signaling pathway is dramatically impaired in the *arfa^sept^* lines. Indeed, colonies from both *arfa^sept^* lines largely phenocopy *afb1,2,3,4* and *iaa2mDII* colonies in which the auxin signaling pathway is severely inhibited. We sought to assess the degree to which the auxin response is hindered in each mutant line compared to wild type. To do so, we grew each line on medium supplemented with 5.0 µM of the synthetic auxin 1-Naphthaleneacetic acid (NAA). Wild-type plants grown on NAA are chlorotic and develop few leafy gametophores (Bascom et al. 2023; Lavy et al. 2016; Plavskin et al. 2016). Therefore, we assessed auxin signaling in each of these mutants by measuring the chlorophyll content and gametophore number of each line when grown with or without NAA supplementation (Figure 1). After growing for 21-days on 5.0 µM NAA, wild-type tissue had 30% of the chlorophyll content as the mock treated plants (Figure 1B). In contrast, chlorophyll content in the *afb1,2,3,4, iaa2mDII,* and *arfa^sept^* lines was only slightly affected by 5.0 µM NAA treatment (Figure 1B). These slight differences were not significant (paired T-Test, p>0.05), except for the *iaa2mDII* line. The *iaa2mDII* line may be slightly less NAA-resistant than the others because it retains two wild-type *Aux/IAA* genes. This could further explain why the *iaa2mDII* phenotype is not as severe as *afb1,2,3,4.* Therefore, the *iaa2mDII* line can mildly respond to prolonged NAA treatment.

In agreement with previous reports, wild type produces an average of 25.9 gametophores/colony on control medium, yet only 0.2 gametophores/colony on medium supplemented with 5.0 µM NAA after 21 days (Figure 1C). The *afb1,2,3,4* and *iaa2mDII* lines never developed gametophores (Figure 1C). These results corroborate earlier findings in which other “strong” *PpAux/IAA* degron mutations result in lines that failed to initiate gametophores (Prigge et al. 2010; Tao and Estelle 2018). (Prigge et al. 2010) demonstrated that *PpAFB* RNAi knock-down plants lack recognizable caulonemata. Therefore, it was not surprising that the *afb1,2,3,4* line failed to develop gametophores. Given the severity of both the *iaa2mDII* and *afb1,2,3,4* lines, we were surprised that *arfa^sept^* colonies were capable of producing gametophores. Indeed, both *arfa^sept^* lines produced 1-3 gametophores/colony, independent of NAA supplementation (Figure 1C). These results suggest that, while auxin signaling plays a significant role in gametophore development, the induction of auxin sensitive genes in response to auxin is not essential for gametophore initiation and development.

### Degree of auxin signaling inhibition has varied effects on protonemal development

Having observed differences in gametophore production among *afb1,2,3,4, iaa2mDII,* and *arfa^sept^* mutants, we then asked if there were developmental defects with these mutants before gametophore initiation. Prior to producing gametophores, *P. patens* grows as a filamentous tip-growing tissue collectively referred to as protonemata. Moss protonemata are comprised of two developmentally sequential cell types– chloronemata and caulonemata. An individual chloronemal apical cell undergoes a cell differentiation event to become a caulonemal cell. Previous studies indicate that this transition is an auxin-dependent event (Plavskin et al. 2016; Prigge et al. 2010; Prigge and Bezanilla 2010; Thelander, Landberg and Sundberg 2018). Of the many differences between the two cell types, a clearly distinguishable characteristic is that chloronemata have cell cross walls that are perpendicular to the axis of growth, while caulonemata have oblique cell cross walls. Therefore, to characterize the chloronema to caulonema transition, we measured all clearly discernable cross wall angles, relative to the axis of growth of the protonemal filament, in seven-day-old plants regenerated from single protoplasts (Figure 2). Because the plants start from identically sized single cells, they are nearly synchronized. After seven days of growth many filaments have undergone the chloronema-caulonema transition such that the average cross wall angle was 111.7°±14.0° across more than 600 cells per line (Figure 2B). The data formed a bimodal distribution around that average, indicative of the two cell types. Previous work used 110° as the demarcation between chloronemata from caulonemata (Bascom et al. 2023), and we utilized the same rule here. We found that 43.8% of wild-type cells were chloronemal and 56.2% were caulonemal (Figure 2C). In stark contrast, the *afb1,2,3,4* mutant had an average cross wall angle of 100.2°±11.1° and no bimodal distribution (Figure 2B). Further, 81.4% of *afb1,2,3,4* cells were chloronemata and only 18.6% were caulonemata (Figure 2C). Interestingly, the *iaa2mDII* and *arfa^sept^* lines had similar cross wall averages (∼107°±14°) and cell type composition (57-60% chloronemata, 40-43% caulonemata) (Figure 2B,C). These results demonstrate that the loss of AFB function has more severe consequences to growth and development than the *iaa2mDII.* Further, the transcriptional repression exerted by the *iaa2mDII* mutation is enough to fully inhibit gametophore development, but only partially inhibit protonemal development.

**Figure 2.**
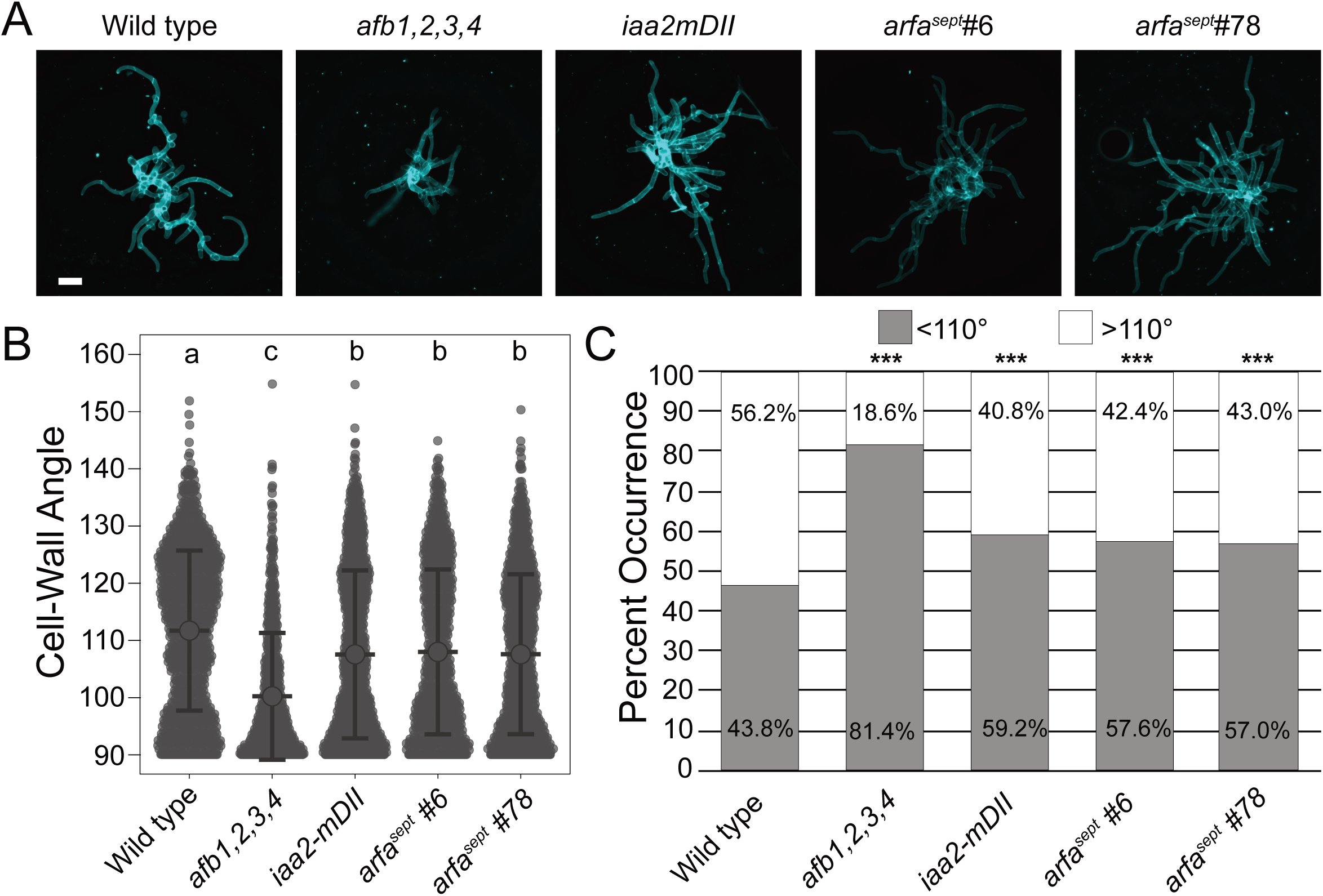
Protonemal development is delayed in auxin signaling mutants. A) Representative micrographs of seven-day-old plants regenerated from single protoplasts. Cell walls stained with calcofluor. Scale bar is 100 µm. B) Protonemal cell wall angles of seven-day-old plants regenerated from single protoplasts, grown on BCDAT. *n* ≥ 679 cells walls per line across three replicates. Large circle and error bars are average and standard error, respectively. Letters are the result of Tukey HSD post-hoc test of an ANOVA, α=0.01. C) Proportion of cell wall angles that fall above (white) or below (grey) 110° relative to the angle of filament growth. *** = p<0.001, Kolmogorov– Smirnov test.

Conversely, in the absence of auxin-response gene activation by the ARFas, *arfa^sept^* plants can still produce both gametophores and caulonemata, but at diminished rates compared to wild type.

### arfa^sept^ mutations alter basal expression of auxin response genes

The results of the gametophore production assay suggest that auxin signaling in the *arfa^sept^* mutants is not as inhibited as plants with complete (in the case of *afb1,2,3,4)* or severe (*iaa2mDII)* repression of auxin response genes. Yet, measuring the chloronema-to-caulonema transition suggests that auxin signaling is similarly inhibited in *iaa2mDII* and *arfa^sept^* lines. To test this directly, we measured mock and auxin induced expression of four auxin response genes (Figure 3). To facilitate auxin induction, we exposed seven-day-old regenerated homogenized tissue to 5 µM IAA for two hours in liquid medium. To quantify the transcriptomic auxin response, we measured the expression of four genes which previous reports found to be auxin-regulated (Lavy et al. 2016; Prigge et al. 2010). Consistent with those findings, auxin treatment produced a strong induction in each gene (Figure 3). Strikingly, auxin induction was nearly completely abolished in all mutants. While *ZHL* expression in *afb1,2,3,4* and *arfa^sept^ #6* was statistically higher after auxin treatment, the fold-change (1.75x and 2.44x, respectively) is far less than what occurs in wild type (30.26x) (Figure 3C). Interestingly, the expression of each auxin-response gene in mock treated tissue was substantially below that of wild type in the *afb1,2,3,4* and *iaa2mDII* mutants. This reduction of basal gene expression supports a model in which stabilized Aux/IAA protein further represses the expression of auxin response genes. Further, when detectable, the expression level of each target gene was lower in *afb1,2,3,4* than *iaa2mDII*. Meanwhile, gene expression in mock treated tissue was much closer to wild type levels in each *arfa^sept^* line. These data suggest that auxin response genes are not repressed *per se* in *arfa^sept^*lines, but rather simply cannot be induced. Together, these data highlight that complete (*afb1,2,3,4*) and severe (*iaa2mDII*) transcriptomic repression have more dramatic effects on gene expression than the absence of genetic activators (*arfa^sept^)* (Figure 4). *PpIAA2* expression in *arfa^sept^* lines was a notable exception to the general trend observed with other genes (Figure 3D). Indeed, *PpIAA2* expression was nearly two-fold higher in mock treated *arfa^sept^* than in mock treated wild type. This observation highlights the presence of complex feed-back loops among auxin signaling genes. In *arfa^sept^,* the net result is presumably more PpIAA2 transcriptional repressor. Though, without ARFa proteins through which PpIAA2 protein would affect gene repression, it is unclear if the measured increase in *PpIAA2* expression has a significant impact on total auxin response gene expression in the *arfa^sept^* lines.

**Figure 3.**
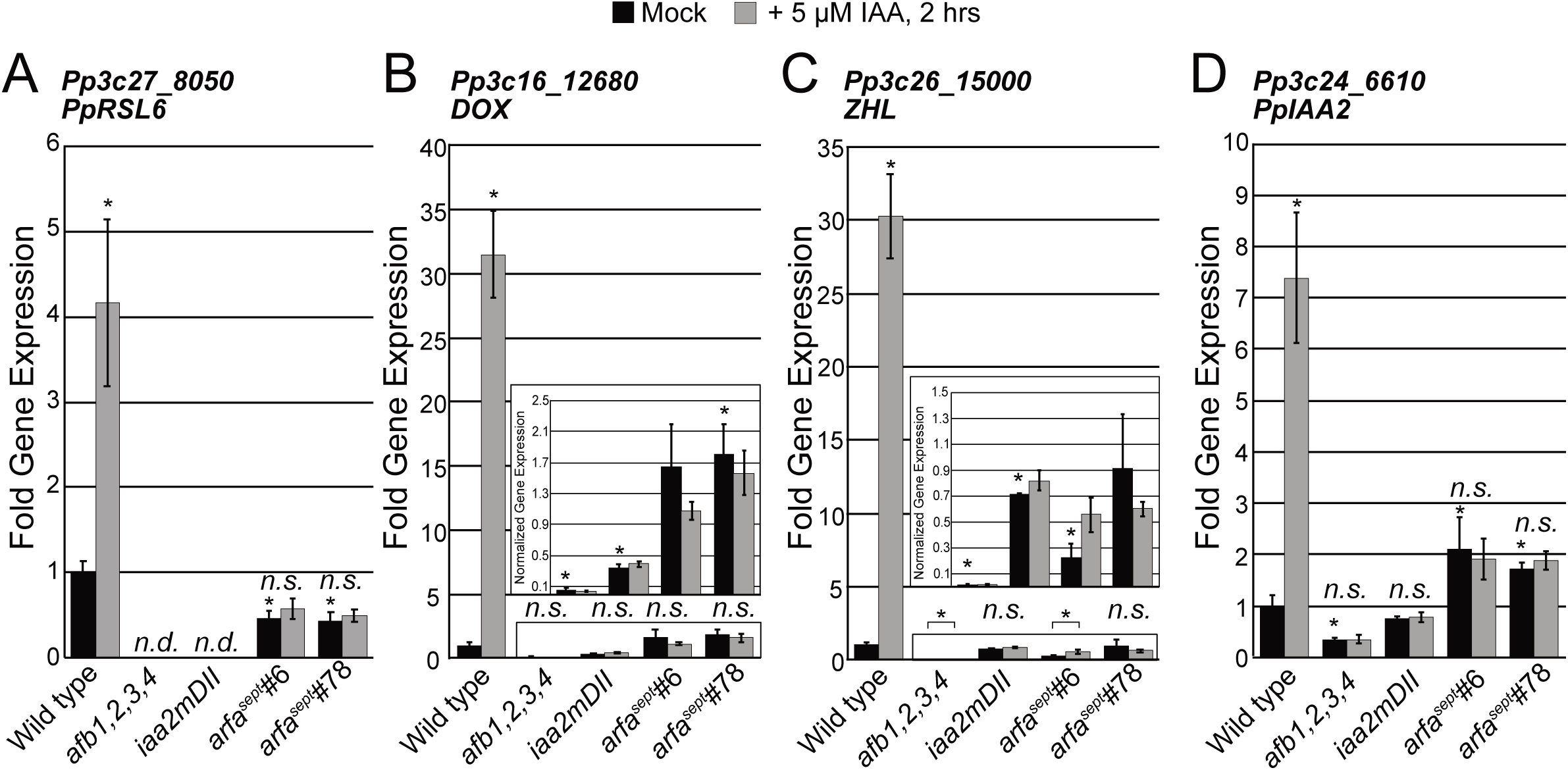
Fold gene expression change upon two hours of 5.0 µM auxin treatment. A) *PpRSL6,* B) *PpDOX*, C) *PpZHL,* and D) *PpIAA2* expression is up-regulated in wild type upon auxin treatment. Student’s T-Test of normalized gene expression between lines compares expression to mock-treated wild type. Within a mutant line, fold-change significance was compared to mock-treated expression levels. * indicates p<0.05, Student’s T-Test, three biological replicates with two technical replicates each.

**Figure 4.**
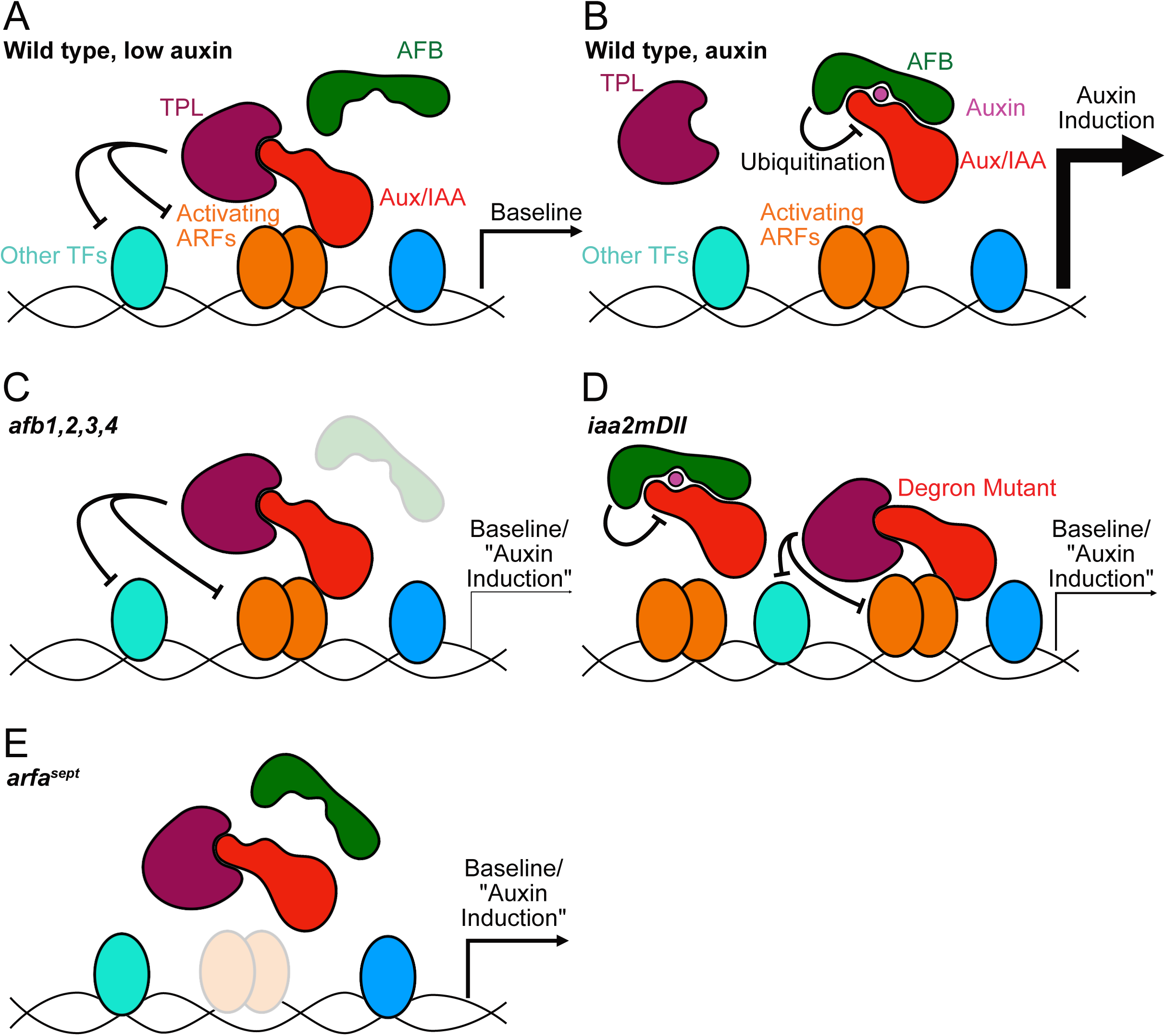
Model of auxin de-repression. When the auxin concentration in the nucleus is low (A), the effect of ARFas, as well as other non-ARF transcription factors, is limited by Aux/IAA-mediated recruitment of the TPL co-repressor closing the locus. As auxin levels rise (B), AFBs ubiquitinate Aux/IAAs, thereby allowing all transcription factors to act on the locus. As a result, there is a significant difference in gene expression between baseline and auxin induction. In *afb1,2,3,4* mutants (C) Aux/IAAs are not degraded, and therefore the promoter remains closed. In the *iaa2mDII* degron mutant (D), one of three Aux/IAAs is stabilized, thereby resulting in only partial promoter closure. In *arfa^sept^* lines, Aux/IAAs are not recruited to the promoter regions. While gene expression remains diminished without activating ARFs, other non-ARF transcription factors are allowed to provide baseline gene expression. However, there is no auxin mediated induction.

## Discussion

Clade A activating ARFs are the best characterized of the ARFs and their role in the nuclear auxin signaling pathway is well understood (Wójcikowska, Belaidi and Robert 2023). In *Arabidopsis,* 5 of the 23 ARFs are activating ARFs (Cancé et al. 2022). Activating ARFs in *Arabidopsis* have different roles in development. *MP/ARF5* functions during embryogenesis and later during vascular patterning in leaves (Hardtke and Berleth 1998). Likewise, *AtARF7* and *AtARF19* are each involved in leaf growth and lateral root initiation (Okushima et al. 2005), while *AtARF6* and *AtARF8* function during flower development (Nagpal et al. 2005).Complete repression via mutating all *TIR1/AFB* genes is embryo lethal (Prigge et al. 2020), and severe repression via *Aux/IAA* degron mutants can dramatically hinder growth (Luo, Zhou and Zhang 2018; Rinaldi et al. 2012; Tian, Uhlir and Reed 2002). Given the severity of activating ARF recessive phenotypes, and the overlapping function of the activating ARFs, it has been technically challenging to conduct detailed investigations of growth and development the absence of auxin-response gene induction in *Arabidopsis*.

To understand plant development in the absence of auxin-mediated gene activation, previous work focused on the bryophyte *M. polymorpha* because all auxin-mediated gene activation occurs via the single activating ARF-*MpARF1* (Kato et al. 2017). What remained unclear, however, is whether or not the absence of auxin-mediated gene activation has similar consequences to growth as the constitutive repression of auxin response genes. Here, we present the second model organism, *Physcomitrium patens*, in which all clade A ARFs have been mutated (*arfa^sept^).* Further, we present evidence that there is a clear transcriptomic and developmental difference between the absence of auxin-mediated gene activation and constitutive repression (Figure 4).

Superficially, the *arfa^sept^*, *afb1,2,3,4,* and *iaa2mDII* colonies are phenotypically similar. Each exemplifies the well-described auxin insensitivity phenotypes in *P. patens*– small and round. Other researchers have observed this phenotype when treating *P. patens* with the auxin biosynthesis inhibitor L-Kyn (Lavy et al. 2016; Nemec□Venza et al. 2022), or expressing a miRNA-insensitive clade B repressing ARF (Plavskin et al. 2016). Recent studies into another clade of activating ARF, the clade D ARFs, found that *P. patens* plants carrying null lesions of both *ARFds* were also more round than wild type (Bascom et al. 2023). This has generally been attributed to the lack of spreading caulonemal filaments, the development of which is an auxin-mediated process (Eklund et al. 2010; Johri and Desai 1973; Pressel, Ligrone and Duckett 2008).

After 21-days of growth, however, there were apparent differences between the three sets of mutants. Indeed, *afb1,2,3,4* colonies were typically smaller than the other lines. We also noted this trend in seven-day-old regenerated protoplasts, suggesting the *afb1,2,3,4* plants grow more slowly than the other lines. While gametophore production was completely abolished in both *afb1,2,3,4* and *iaa2mDII,* the *arfa^sept^* colonies were able to produce a few gametophores. The lack of gametophore production in the mutants is stark compared to wild type. Therefore, it is tempting to categorize the *arfa^sept^*lines with the other mutants. Indeed, all mutants are similarly insensitive to NAA-induced chlorosis. Yet, the fact that *arfa^sept^* lines could make gametophores at all was counter to our expectations. Whether these gametophores are wild type in morphology will have to be studied at a later date. Additionally, these mutants will allow further dissection of auxin’s role in the switch between two-dimensional (spreading filamentous protonemata) and three-dimensional (gametophores) growth as our results demonstrate that auxin-mediated gene activation is not strictly necessary (Moody 2022; Nemec□Venza et al. 2022; Raquid, Jaeger and Moody 2023).

Based on the results of the colony experiments, we sought to describe the *arfa^sept^* mutants earlier in development. Specifically, we investigated whether the auxin-mediated chloronema-caulonema transition is similarly affected across the mutants. Much like gametophore development, the chloronema-caulonema transition is an auxin-mediated process (Eklund et al. 2010; Johri and Desai 1973; Pressel, Ligrone and Duckett 2008; Prigge and Bezanilla 2010; Thelander, Landberg and Sundberg 2018). Unsurprisingly, the chloronema-caulonema transition is compromised in *afb1,2,3,4* plants. This finding complements our results that *afb1,2,3,4* colonies do not produce gametophores, in line with complete repression of auxin-mediated developmental processes (Figure 4). Meanwhile, *iaa2mDII* and *arfa^sept^* seven-day-old plants have a reduced chloronema:caulonema ratio compared to wild type, indicative of a delay in development. Surprisingly, the chloronema:caulonema ratio in *iaa2mDII* and *arfa^sept^* seven-day-old plants were nearly identical. One interpretation is that the over-all level of auxin signaling is reduced in *iaa2mDII* to a level similar to the *arfa^sept^* line. In this case, constitutive-repression and the absence of auxin-mediated gene activation have a similar net result (Figure 4). Yet, expression analysis of a few known auxin-regulated genes in these lines indicates that auxin signaling in *iaa2mDII* plants is reduced compared to *arfa^sept^* plants. An alternative interpretation is that specific genes directly responsible in the chloronema-caulonema transition, such as *SHORTINTERNODES (SHI/STY)* (Eklund et al. 2010) and *CLAVATA3/ESR (CLE)* (Nemec□Venza et al. 2022), could be sufficiently expressed in *iaa2mDII* to generate a chloronema:caulonema ratio similar to *arfa^sept^.* While *iaa2mDII* and *arfa^sept^* plants undergo the chloronema to caulonema transition to a similar degree, *iaa2mDII* plants never produce gametophores in our experiments while the *arfa^sept^* plants are capable of producing a few gametophores. Presumably, the disparity is due to auxin-regulated genes that are differentially responsible for either caulonema or gametophore development. Likewise, some genes responsible for gametophore initiation and development have sufficient expression in *arfa^sept^* to result in a few gametophores. Future efforts will have to conduct transcriptome analysis in these mutants to elucidate the whole-transcriptome effects of constitutive repression versus the absence of auxin-mediated gene activation (Figure 4). The work presented here suggests that non-ARF transcription factors on the promoters of auxin response genes provide a basal level of gene expression. Therefore, transcriptomic analysis experiments could identify potentially novel genes and pathways involved in plant development in both auxin dependent and auxin independent ways. Further, the *arfa^sept^* lines are the first plants in which auxin-mediated gene activation has been abolished in an organism with multiple clade A ARFs. These mutants are poised to serve as tools to dissect potential subfunctionalization among the ARFas, as well as cross-species or synthetic biology complementation studies.

## Materials and Methods

### Plant tissue and growth conditions

All growth assays, tissue propagation, and transgenic line generation were conducted in a growth chamber (Percival Scientific, Perry, Iowa, USA) under continuous light, at 25°C. Tissue was grown aseptically on sterile BCDAT medium, as has been described previously (Cove et al. 2009). The Reute ecotype served as wild type, and the background for *arfa^sept^, afb1,2,3,4* and *iaa2mDII* lines (Hiss et al. 2017). The *afb1,2,3,4* and *iaa2mDII* lines were described previously (Bascom et al. 2023). Transformations to generate the *arfa^sept^* were performed in a PEG-mediated manner described previously (Schaefer et al. 1991) with modifications as described in (Bascom et al. 2023).

### Protospacer design, cloning, and *arfa^sept^* mutant generation

To generate the *arfa^sept^* line, we used a CRISPR/Cas9 plasmid system described previously (Mallett et al. 2019). We avoided using the pMZ-Gate Cas9 plasmid, which confers transient Zeocin resistance to transformants, due to the DNA damage Zeocin can cause (Čížková et al. 2019). Therefore, we used CRISPR/Cas9 plasmids with either Hygromycin (pMH-Cas9) or G418/Kanamycin (pMK-Cas9) resistance. We used the web-based CRISPOR tool to design protospacers (Concordet and Haeussler 2018). Initially, we designed sgRNAs to attack two sites in the 5′ end of genes to maximize the chances of generating a frame-shift mutation. The location of all protospacers is indicated in Supplemental Figure 2. For *ARFa5,* we could not recover any mutations with our initial pMH-ARFa5_Cas9 plasmid (protospacers not shown) and we thus re-designed the protospacers to knock out the majority of the coding region with pMK-ARFa5_Cas9. Similarly, pMK-ARFa7_Cas9 resulted in an in-frame *arfa7-6* mutation. Therefore, we generated pMH-ARFa7_FullKO_Cas9 with protospacers indicated in Supplemental Figure 2. To avoid large-scale genome damage, we elected to transform ARFa-targeting CRISPR/Cas9 plasmids alone or in pairs. Recovered lines were genotyped by PCR amplification of the locus before an individual mutant was transformed with the next plasmids. We co-transformed the *arfa8* mutant with pMH-ARFa6_Cas9 and pMK-ARFa7_Cas9. We then co-transformed the triple *arfa6,7,8* mutant with pMK-ARFa5_Cas9 and pMH-ARFa4_Cas9. We then co-transformed the quintuple *arfa4,5,6,7,8* quintuple mutant with pMH-ARFa1_Cas9 and pMK-ARFa3_Cas9 and isolated two *arfa^sept^* mutant lines– *arfa^sept^#6 and #26*. As mentioned above, these lines carried an in-frame deletion of *ARFa7,* and so we transformed *arfa^sept^#26* to with pMH-ARFa7_FullKO_Cas9 to generate *arfa^sept^#78*.

### Chlorophyll Extraction

Chlorophyll extractions were performed as described previously (Bascom et al. 2023). Whole 21-day-old colonies were removed from the BDCAT medium with tweezers, lightly dried with paper towels, weighed, and placed in a 2.0 mL Eppendorf microcentrifuge tube with 1.0 mL of Methanol. The chlorophyll extraction incubated over-night in the dark at laboratory room temperature. Absorbances of 652 nm, 665 nm, and 750 nm were measured via SmartSpec Plus spectrophotometer.

### Microscopy and Image Analysis

Images of 21-day-old colonies were acquired with a Nikon SMZ1500 microscope via a DS-Ri1 color camera. Seven-day-old regenerated protoplasts, treated briefly with Fluorescent Brightener 28 (Calcofluor-white, Sigma) to stain the cell walls, were imaged with a BZ-X810 Keyence benchtop microscope. Micrographs were opened with ImageJ to manually measure cross wall angles.

### Gene Expression Analysis

Seven-day-old tissue, regenerated from light homogenization, was harvested aseptically into 50 mL conical tubes with 20 mL of filter-sterilized liquid BCD medium (BCDAT minus ammonium tartrate) supplemented with 5.0 µM IAA or solvent control. Tissue was then filtered out of the treatment solution, lightly dried with paper towels, and flash-frozen with liquid nitrogen until RNA extraction. RNA extraction was performed with the Zymo Quick-RNA Plant Miniprep kits (Zymo Research, Irvine, CA, USA) according to the manufacturer’s instructions. cDNA synthesis was performed via the Maxima H Minus cDNA Synthesis Master Mix (Thermo Fischer Scientific, Waltham, MA, USA). Quantitative PCR experiments used lab-made reagents, with measurements made on a CFX Connect Real Time System thermocycler/plate reader (BioRad). We used *ADENINE PHOSPHORIBOSYLTRANSFERASE* (*APT/ Pp3c8_16590V3.1*) as a housekeeping gene, as described previously (Bail, Scholz and Kost 2013). We chose auxin-response genes based on auxin induction described previously (Lavy et al. 2016). For the expression meta-analysis of ARFas, *IAA2*, and the AFBs, we extracted reported expression levels from three publicly available datasets (Lavy et al. 2016; Ortiz-Ramírez et al. 2016; Perroud et al. 2018). For the Lavy et al, 2016 dataset, we mapped reads using SAMtools (Li et al. 2009) to the version 3 of the *P. patens* genome (Lang et al. 2018). We then calculated Fragments per Kilobase Mapped (FPKM) via the following equation:

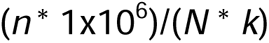

Where *n* is the number of fragments mapped to the region of interest, *k* is the transcript length (in kilobases) and *N* is the total number of mapped fragments. We then normalized the expression of each to the expression of *PpAPT* in protonemata before averaging.

### Data Preparation

Averages, standard deviations, standard error of the mean, and Student’s T-Tests were calculated in Excel (Microsoft). Likewise, bar and 100% stacked column graphs were made in Excel, while dot plots were generated with an R-based web tool (Goedhart 2021). Kolmogorov–Smirnov, ANOVA, and TukeyHSD tests were done in R. Figures were prepared with Affinity Designer (Serif Europe).

---

**Supplemental Figure 1** Gene diagrams of the activating ARFs in *P. patens*, and the mutations obtained via CRISPR/Cas9. Black text denotes original genomic sequence, green text indicates the start codon of *ARFa3*. Red text in *ARFa5* is the premature stop codon. Pink text highlights insertions, backwards text indicates an inverted insert (when the inserted sequence could be identified). sgRNA targets are indicated with red arrows, while the sequence is highlighted with a red bar. The yellow sgRNAs shown in *ARFa7* are where the protospacers were designed to mutate *arfa7-6*.

**Supplemental Figure 2** Predicted peptide sequences resulting from CRISPR/Cas9 mutations to ARFas. Bold text indicates where the peptide sequence diverges from the wild type sequence.

**Supplemental Figure 3** Relative expression level and pattern of select auxin-signaling genes vary across P. patens *developmental stages.* Expression of *PpARFas*, *PpIAA2* and *PpAFBs,* relative to *PpAPT* expression in protonemata from (A) Ortiz-RamÌrez et al., 2015, (B) Perroud et al., 2018, and (C) Lavy et al., 2016. Where necessary, data was aggregated from developmental sub-stages (e.g. “green sporophyte” and “brown sporophyte” to “sporophyte”) by simple average. Error bars are standard error.

## Notes

### Competing Interest Statement

The authors have declared no competing interest.

