## Supplementary figures and images for "The role of auxin-mediated gene activation in the bryophyte, *Physcomitrium patens*"

### Supplemental Figure 1

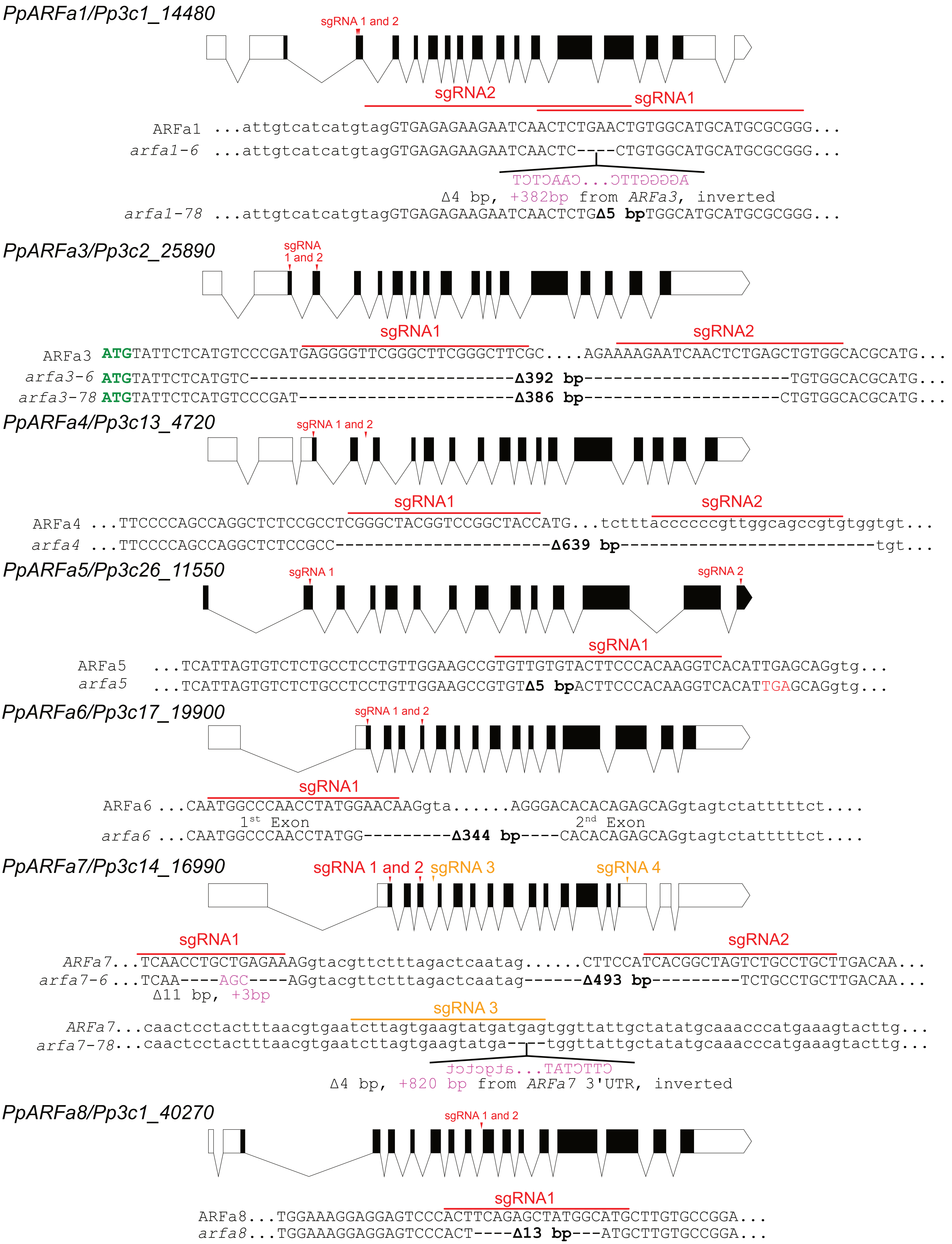

### Supplemental Figure 3

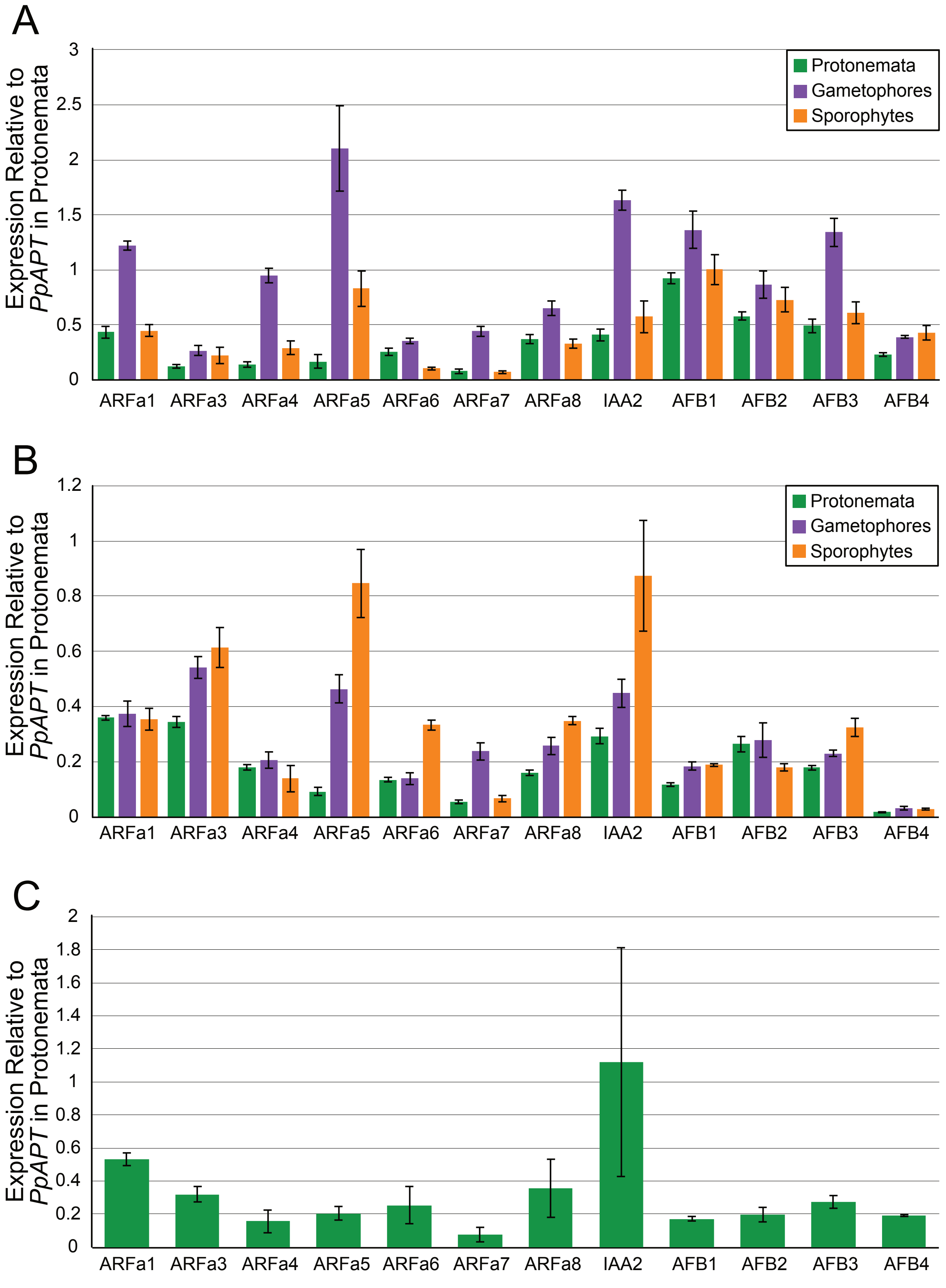
