## Supplemental Figure 2 for "The role of auxin-mediated gene activation in the bryophyte, *Physcomitrium patens*"

ARFa1 MYSHASMRGSGYGPPAMDQGERRINSELWHACAGPLVSLPPVGSQVVYFPQGHSEQVAVSTQKEADTHIPNYPNLRPHLVCTLDNITLHADLETDE...
arfa1-6 MYSHASMRGSGYGPPAMDQGERRINSEL**ILFSPWTIVEEAKPEPLCGMHARGHWFHFLQWAAKWFTFHKVTLSRLLCQPKRKLTLIFPTTQIFDHT***
arfa1-78 MYSHASMRGSGYGPPAMDQGERRINS**VACMRGAIGFTSSSGQPSGLLSTRSL***

ARFa3 MYSHVPMRGSGFASSTIVQGEKRINSELWHACAGPLVSLPPVGSQVVYFPQGHSEQVAVSTQKEADIHIPNYPNLRPHLICTLENVTLHADLETDD...
arfa3-6 MYSHV--**CGTHVRDHWFHFLLWVAKWFTSHKVTLSRLQCQLKRKLTFIFQTTQTFDHTSSVHLKMSPFMRILKRTMFMHRWFLFQRKIPKKKRCCYQM***
arfa3-78 MYSHVP**ICGTHVRDHWFHFLLWVAKWFTSHKVTLSRLQCQLKRKLTFIFQTTQTFDHTSSVHLKMSPFMRILKRTMFMHRWFLFQRKIPKKKRCCYQM***

ARFa4 MYSSSPARLSASGYGPATMDPGLSLSLSLSLSRSLSYCMCVSEKYIYIYILTHTHTHTHVAASTQKDADAHIPNYPSLPSKIICLLDNVTLHADPE...
arfa4 MYSSSPARLSASG**CCINTEGC***

ARFa5 MYSNPPARLSASGHVSSTMDPGGRRSLNSELWHACAGSLVSLPPVGSRVVYFPQGHIEQVAASTQKEADVPIPNYPSLPSRLFCLLDNVSLHADHE...
arfa5 MYSNPPARLSASGHVSSTMDPGGRRSLNSELWHACAGSLVSLPPVGSRV**LPTRSH***

ARFa6 MYLSSERPPSLGPIMTSMAQPMEQVERRSLNSELWHACAGPLVSLPPVGSRVVYFPQGHTEQVAASTQKEADAHIPNYPNLPSRLVCLLDNVTLHA...
arfa6 MYLSSERPPSLGPIMTSMAQPM**AHRAGCSINAEGGRCSYPQLSQPSITACLSA***

ARFa7 MYLCNERLPSVGPSVGSMAQPAEKVEKRSLNSELWHACAGPLVLLPPVGSRVVYFPQGHTEQVAASMQKEVDAHIPNYPNLPSRLVCLLDNVTLHADIE...
arfa7-6 MYLCNERLPSVGPSVGSMAQ**QQ**-------------------------------------------------------------------------ADIE...
arfa7-78 MYLCNERLPSVGPSVGSMAQ**QQSACLTMSRFMLILRRMKCMLR***

ARFa8 MEVVAEEESRPCGATVWSGFLRVERRSPTSELWHACAGPLVSLPPIGSRVVYFPQGHTEQVAASTQREAETHIPNYPSLPSRLVCLLDNVTLHADLET...
arfa8 MEVVAEEESRPCGATVWSGFLRVERRSPT**MACLCRTLGLTSSYWQPGGVLSSRSYGTGSSINSKRG***
